# The transcriptional and splicing changes caused by hybridization can be globally recovered by genome doubling during allopolyploidization

**DOI:** 10.1101/2020.09.06.285262

**Authors:** Jinxia Qin, Ruirui Mo, Hongxia Li, Zhongfu Ni, Qixin Sun, Zhenshan Liu

**Affiliations:** State Key Laboratory of Crop Stress Biology for Arid Areas, College of Agronomy; Northwest A&F University; Yangling, Shaanxi, 712100; China; State Key Laboratory for Agrobiotechnology, Key Laboratory of Crop Heterosis and Utilization (MOE), Beijing Key Laboratory of Crop Genetic Improvement; China Agricultural University; Beijing, 100193; China; Department of Cell and Developmental Biology; John Innes Centre; Norwich Research Park, Norwich, NR4 7UH; United Kingdom

**Keywords:** Allopolyploidization, Heterosis, Splicing, Transcriptome, Wheat, Brassica

## Abstract

Allopolyploidization, which involves hybridization and genome doubling, is a key driving force in higher plant evolution. The transcriptome reprogramming that accompanies allopolyploidization can cause extensive phenotypic variations, and thus confers allopolyploids higher evolutionary potential than their diploid progenitors. Despite many studies, little is known about the interplay between hybridization and genome doubling in transcriptome reprogramming during allopolyploidization. Here, we performed genome-wide analyses of gene expression and splicing changes during allopolyploidization in wheat and brassica lineages. Our results indicated that both hybridization and genome doubling can induce genome-wide transcriptional and splicing changes. Notably, the gene transcriptional and splicing changes caused by hybridization can be largely recovered to parental levels by genome doubling in allopolyploids. Since transcriptome reprogramming is an important contributor to heterosis, our results revealed that only part of the heterosis in hybrids can be fixed in allopolyploids through genome doubling. Therefore, our findings update the current understanding of the permanent fixation of heterosis in hybrids through genome doubling. In addition, our results indicated that a large proportion of the transcriptome reprogramming in interspecific hybrids was not caused by the merging of two parental genomes, providing novel insights into the mechanism of heterosis.

## Introduction

Allopolyploidization, which involves interspecific hybridization and genome doubling, is a critical driving force in plant evolution and crop domestication (Alix et al., 2017; Baduel et al., 2018; Chen, 2013; Guo and Han, 2014; Monnahan et al., 2019). The transcriptome reprogramming that accompanies allopolyploidization can cause various phenotypic novelties, and thus confers higher evolutionary potential and plasticity in allopolyploids (Alix et al., 2017; Hegarty et al., 2008; te Beest et al., 2012; Wang et al., 2006; Yoo et al., 2013; Zhao et al., 2013). The interplay between hybridization and genome doubling in composing the transcriptome reprogramming in allopolyploids is a mechanistic basis underlying allopolyploidization but is still elusive (Qiu et al., 2020).

Though the transcriptional changes in allopolyploids compared with their progenitors have been frequently reported (Chen, 2013; Hegarty et al., 2008; Wang et al., 2006; Xu et al., 2014; Yoo et al., 2013; Zhao et al., 2013), very few studies have dissected the respective effects of hybridization and genome doubling during allopolyploidization (Hao et al., 2017; Hegarty et al., 2006). A previous study in *S. cambrensis* using ‘anonymous’ cDNA microarrays found that the expression changes in many floral genes which resulted from hybridization were attenuated after genome doubling (Hegarty et al., 2006). However, the inherent limitations of ‘anonymous’ cDNA microarrays make it difficult to distinguish the expression of homoeologous genes from different subgenomes, which compromises their conclusion (Zhao et al., 2014). A recent RNA-Seq study in wheat reported that hybridization mainly caused the down-regulation of genes from the D subgenome, which can be partially restored by genome doubling (Hao et al., 2017). However, very few genes in the A and B subgenomes were found to be affected during allopolyploidization (Hao et al., 2017). To date, little is known about to what extent the hybridization-induced transcriptional changes can be affected by genome doubling.

Splicing regulation is a key post-transcriptional regulatory mechanism in eukaryotes (Syed et al., 2012). To our knowledge, only few studies have identified individual genes showing splicing changes in allopolyploids (Saminathan et al., 2015; Terashima and Takumi, 2009; Zhou et al., 2011). Thus, how hybridization and genome doubling affect gene splicing at whole-genome scale is still poorly understood.

In this study, we reanalyzed the public available RNA-Seq datasets and examined the transcriptome reprogramming during allopolyploidization in wheat and brassica lineages (Hao et al., 2017; Zhang et al., 2018). We found that both hybridization and genome doubling can induce genome-wide changes at gene transcriptional and splicing levels. Notably, a large proportion of the hybridization-induced transcriptional and splicing changes in hybrids can be recovered to parental levels in allopolyploids after genome doubling. Since transcriptome reprogramming is an important contributor to heterosis (Chen, 2010; Chen, 2013), our results indicate that the heterosis in interspecific hybrids can only be partially fixed in allopolyploids, which updates a longstanding typical theory “heterosis in interspecific hybrids can be permanently fixed through genome doubling” (Chen, 2010; Chen, 2013; Comai, 2005). Additionally, our results indicated that much of the transcriptome reprogramming in interspecific hybrids was not caused by the merging of two parental genomes, providing novel insights into the mechanism of heterosis.

## Results and Discussion

To investigate the effects of hybridization and genome doubling on transcriptome reprogramming during allopolyploidization, we reanalyzed the previously published RNA-Seq datasets of three interspecific crossing combinations in wheat and brassica lineages (Hao et al., 2017; Zhang et al., 2018) (Figure 1A; Supplemental Table 1; Supplemental Figure 1). For wheat combinations, two tetraploids of *T. turgidum* (AABB) were crossed with diploid *Ae. tauschii* (DD) to produce triploid hybrids (ABD) whose genomes were then doubled to generate allohexaploid wheat (AABBDD) (Figure 1A) (Hao et al., 2017). Similarly, diploids *B. rapa* (A_r_A_r_) and *B. oleracea* (C_o_C_o_) were crossed to produce a hybrid (A_r_C_o_) which was used in the generation of allotetraploid brassica (A_r_A_r_C_o_C_o_) (Figure 1A) (Zhang et al., 2018). To examine the respective effects of hybridization and genome doubling on gene expression, significantly differentially expressed genes (DEGs) were identified by comparing hybrids with parents (Hybrid-vs-Parents) and allopolyploids with hybrids (Allopolyploid-vs-Hybrid), with criteria “expression fold change >=2 and false discovery rate (FDR) < 0.05” (Supplemental Tables 2-4). We found that both hybridization and genome doubling can induce dramatic expression changes in thousands of genes (Figure 1B; Supplemental Figure 2). The combined effect of hybridization and genome doubling was further examined by comparing allopolyploids with parents (Allopolyploid-vs-Parents). Notably, for most comparisons (subgenomes A and B in wheat and subgenome A_r_ in brassica), much less DEGs were caused by the combined effect than those caused by hybridization or genome doubling alone (Figure 1B; Supplemental Figure 2). For example, for subgenome A in wheat combination 1, up to 4,759 and 4,735 DEGs were induced by hybridization and genome doubling, respectively, while only 1,849 DEGs were caused by their combined effect (Figure 1B). For subgenome D in wheat and subgenome C_o_ in brassica, the number of DEGs caused by the combined effect was still much less than the sum of DEGs caused by these two events alone (Figure 1B; Supplemental Figure 2).

**Figure 1.**
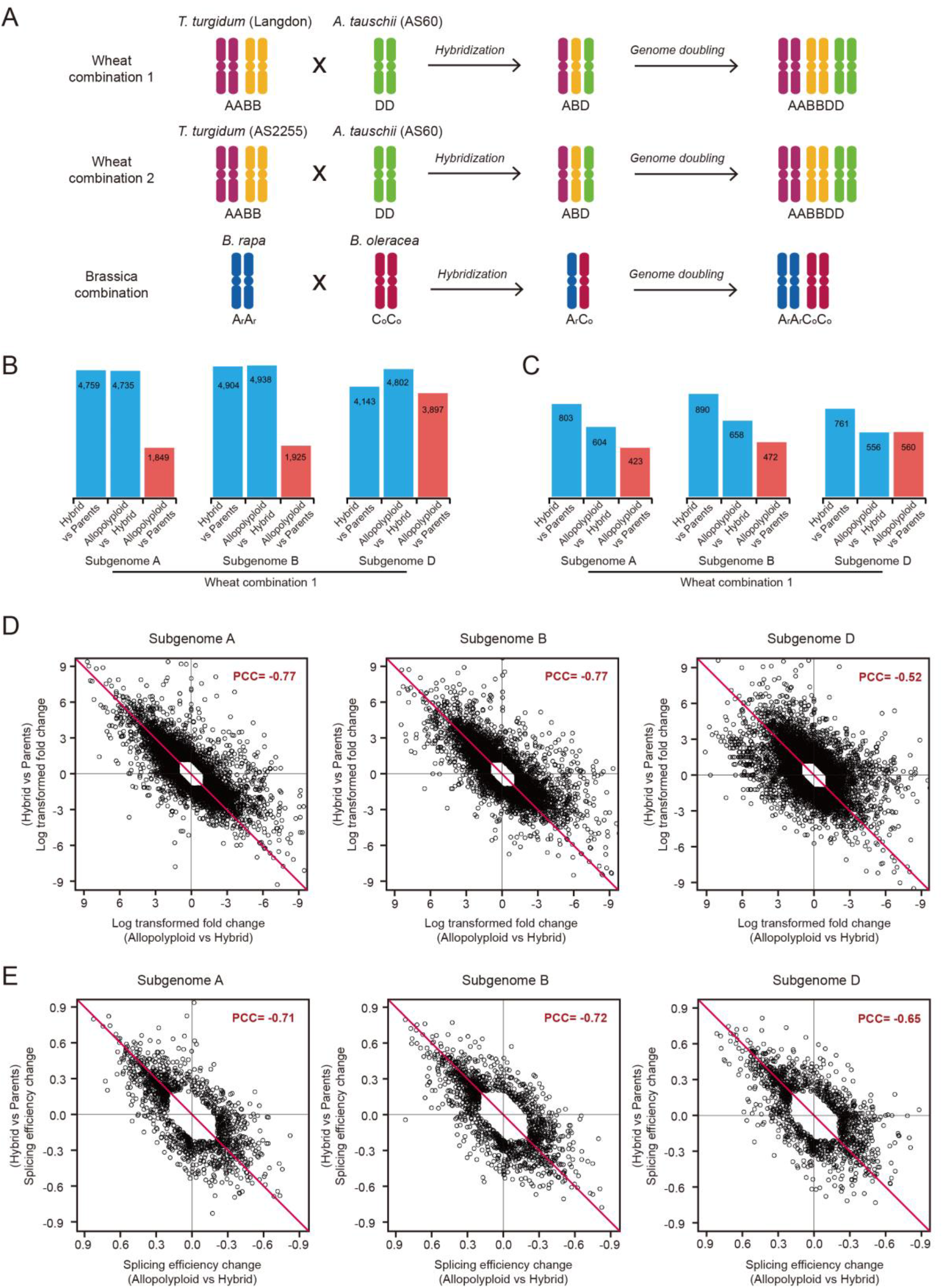
Comparison of gene expression and splicing efficiency changes induced by hybridization and genome doubling during allopolyploidization. **(A)** Schematic representation of the synthesis of allopolyploid wheat and brassica. These hybrids and allopolyploids were produced in previous studies, which provided the datasets used in this study (Hao et al., 2017; Zhang et al., 2018). For wheat combinations, two tetraploids, *T. turgidum* (AABB) ssp. *durum* cv. Langdon (LDN) and ssp. *turgidum* accession AS2255, were crossed with diploid *Ae. tauschii* ssp. *tauschii* accession AS60 (DD) to produce the two triploid hybrids (ABD). The resulting triploids were used to generate the allohexaploid wheat (AABBDD) through genome doubling. For the brassica combination, two diploids *B. rapa* (A_r_A_r_) and *B. orleracea* (C_o_C_o_) were crossed to produce the diploid hybrid (A_r_C_o_), which was then used to generate the allotetraploid brassica (A_r_A_r_C_o_C_o_) through genome doubling. **(B)** The number of differentially expressed genes (DEGs) caused by hybridization, genome doubling and allopolyploidization in wheat combination 1. The numbers of DEGs caused by hybridization (Hybrid-vs-Parents, blue bars), genome doubling (Allopolyploid-vs-Hybrid, blue bars) and allopolyploidization (Allopolyploid-vs-Parents, red bars) are shown for each subgenome. **(C)** The number of differentially spliced introns (DSIs) caused by hybridization, genome doubling and allopolyploidization in wheat combination 1. The numbers of DSIs caused by hybridization (Hybrid-vs-Parents, blue bars), genome doubling (Allopolyploid-vs-Hybrid, blue bars) and allopolyploidization (Allopolyploid-vs-Parents, red bars) are shown for each subgenome. **(D)** Significantly negative correlations between gene expression fold changes caused by hybridization and genome doubling in wheat combination 1. All DEGs identified from Hybrid-vs-Parents, Allopolyploid-vs-Hybrid and Allopolyploid-vs-Parents were analyzed and plotted. The Pearson Correlation Coefficient (PCC) of each comparison is shown (*p* values < 2.2e-16 for all comparisons). **(E)** Significantly negative correlations between splicing efficiency changes caused by hybridization and genome doubling in wheat combination 1. All DSIs identified from Hybrid-vs-Parents, Allopolyploid-vs-Hybrid and Allopolyploid-vs-Parents were analyzed and plotted. The Pearson Correlation Coefficient (PCC) of each comparison is shown (*p* values < 2.2e-16 for all the comparisons).

To examine the genome-wide splicing changes during allopolyploidization, the splicing efficiency of each intron was calculated (see Methods, Supplemental Figure 3). Significantly <u>d</u>ifferentially <u>s</u>pliced <u>i</u>ntrons (DSIs) were identified with criteria “splicing efficiency change >=20% and FDR <0.05” (Supplemental Tables 5-7) (Brooks et al., 2011). We found that both hybridization and genome doubling events can induce genome-wide splicing efficiency changes (Figure 1C; Supplemental Figure 4). For example, in wheat combination 1, 2,454 and 1,818 introns were identified as DSIs caused by hybridization and genome doubling, respectively (Figure 1C). Intriguingly, for most comparisons (subgenomes A and B in wheat and subgenome A_r_ in brassica), there were fewer DSIs caused by the combined effect of hybridization and genome doubling compared to those caused by either event alone (Figure 1C; Supplemental Figure 4). For example, for subgenome A in wheat combination 1, 803 and 604 DSIs were caused by hybridization and genome doubling, respectively, while only 423 DSIs were identified due to their combined effect (Figure 1C). For subgenome D in wheat and subgenome C_o_ in brassica, the DSIs caused by the combined effect were also much less than the total DSIs caused by these two events alone (Figure 1C; Supplemental Figure 4). Collectively, these results indicated that the combined effect of hybridization and genome doubling on gene expression and splicing is far less than the simple sum of their individual effects, which suggests an interplay between hybridization and genome doubling during allopolyploidization.

The relationship between hybridization and genome doubling was further examined by comparing the gene expression and splicing changes induced by these two events (Figure 1D and 1E; Supplemental Figures 5-8). Significantly negative correlations were observed between the expression fold changes induced by these two events for all the subgenomes in all three cross combinations (Figure 1D; Supplemental Figures 5 and 6). For example, the Pearson correlation coefficient (PCC) is −0.52 to −0.77 for different subgenomes of wheat combination 1 (Figure 1D). This result is consistent with a previous study which showed that the abundance of homeologous transcripts disrupted by hybridization can be partially restored by genome doubling (Hao et al., 2017). Furthermore, significantly negative correlations were also observed between the splicing efficiency changes caused by hybridization and genome doubling, as seen in wheat combination 1, which had a PCC of −0.65 to −0.72 (Figure 1E; Supplemental Figures 7 and 8). Taken together, these results indicate that genome doubling can cause global opposite effects compared to hybridization at both gene expression and splicing levels.

To determine how much of the hybridization-induced transcriptome reprogramming in hybrids can be recovered in allopolyploids after genome doubling, DEGs/DSIs identified in the hybrids were further classified into four groups (see Methods): DEGs/DSIs in the hybrids being recovered to parental levels in the allopolyploids (Group 1), DEGs/DSIs in the hybrids being retained in the allopolyploids (Group 2), DEGs/DSIs in the hybrids being reinforced by genome doubling (Group 3) and others (Group 4) (Figure 2A-2D). Most (89-96%) of the DEGs and DSIs can be classified into Groups 1-3 (Figure 2A-2D). Notably, for A and B subgenomes in wheat, most of the hybridization-induced gene expression and splicing efficiency changes in the hybrids can be recovered to parental levels after genome doubling (Group 1, 61-76% for DEGs and 54-75% for DSIs), while fewer were retained in the allopolyploids (Group 2, 15-29% for DEGs and 20-35% for DSIs) (Figure 2A-2D). For subgenome D in wheat, relatively more DEGs/DSIs in the hybrids were retained in the allopolyploids, but still, ∼32% and ∼39% of the DEGs and DSIs in the hybrids were recovered to parental levels by genome doubling, respectively (Figure 2A-2D). Likewise, in the brassica combination, more hybridization-induced DEGs (45-62%) and DSIs (49-63%) were recovered to parental levels compared with those which retained their hybridization-induced changes (31-42% for DEGs, 27-41% for DSIs) in the allotetraploid after genome doubling (Figure 2A-2D). Conclusively, the hybridization-induced transcriptome reprogramming in hybrids can be globally recovered in allopolyploids after genome doubling.

**Figure 2.**
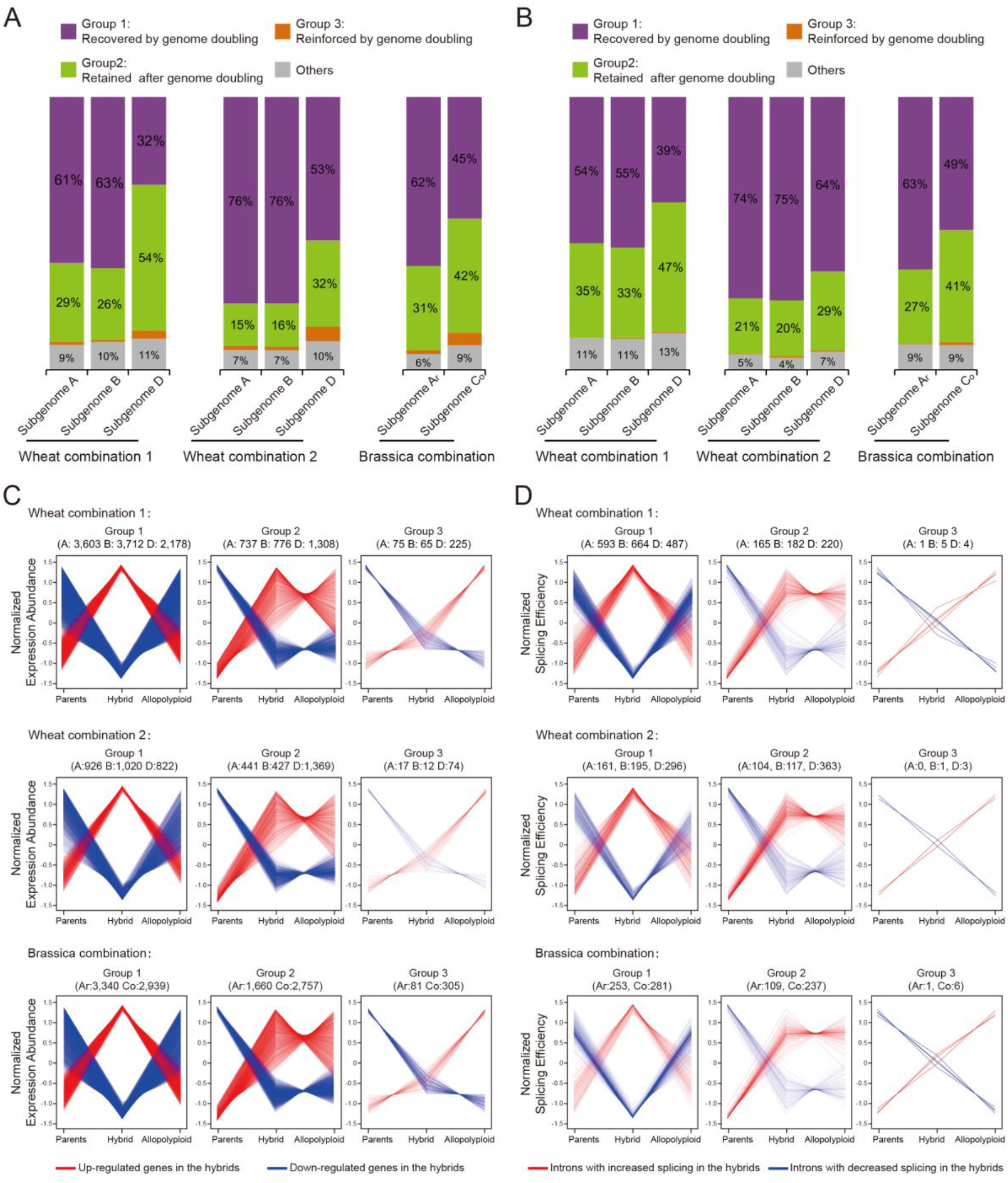
Classification of the differentially expressed genes and differentially spliced introns identified in the hybrids when compared with their parents. **(A)** Classification of DEGs identified in the hybrids compared with their parents. These DEGs were classified into four groups: altered expression in hybrids was recovered to parental level by genome doubling (Group 1), altered expression in hybrids remained in allopolyploid after genome doubling (Group 2), hybridization-induced expression change was reinforced by genome doubling (Group 3) and others (Group 4). **(B)** The expression profiles of the DEGs in Groups 1-3 from the classification analysis in **(A)**. DEGs of the same group from all subgenomes were plotted together and each gene is represented by each single line. The numbers of DEGs in each subgenome are indicated. The up-regulated and down-regulated genes in the hybrids are represented by red and blue lines, respectively. **(C)** Classification of DSIs identified in the hybrids compared with their parents. All DSIs were classified into four groups: altered splicing efficiency in the hybrid was recovered to parental level by genome doubling (Group 1), altered splicing efficiency in hybrids remained in allopolyploids after genome doubling (Group 2), hybridization-induced splicing change was reinforced by genome doubling (Group 3) and others (Group 4). **(D)** The splicing efficiency profiles of the DSIs in Groups 1-3 from the classification analysis in **(C)**. DSIs of the same group from all subgenomes were plotted together and each intron is represented by each single line. The numbers of DSIs in each subgenome are indicated. The introns with increased splicing efficiency and decreased splicing efficiency in the hybrids are represented by red and blue lines, respectively.

Having identified the opposite effects of hybridization and genome doubling, we next attempted to determine their relative contribution to the final transcriptome reprogramming in allopolyploids. The DEGs and DSIs identified from comparisons between the allopolyploids and their parents were further grouped into three clusters according to the relative contribution of hybridization and genome doubling (see Methods): the DEGs/DSIs mainly caused by hybridization (Cluster 1), the DEGs/DSIs mainly caused by genome doubling (Cluster 2), and the DEGs/DSIs significantly contributed by both events with the same trend (Cluster 3) (Figure 3A and 3B; Supplemental Figures 9 and 10). About 34-55% of the gene expression changes (DEGs) and 31-51% of the splicing changes (DSIs) in allopolyploids were found mainly caused by hybridization, and comparable amounts of the DEGs (21-45%) and DSIs (23-43%) were found mainly caused by genome doubling (Figure 3A and 3B; Supplemental Figures 9 and 10). Relatively less DEGs and DSIs were contributed by both events with the same trend (Figure 3A and 3B; Supplemental Figures 9 and 10). These results suggested that both hybridization and genome doubling substantially and comparably contributed to the transcriptome reprogramming in allopolyploids. In addition, the relative contribution of hybridization and genome doubling varied among different subgenomes and species. For example, in wheat combination 1, more DEGs were contributed by hybridization in subgenomes A and B, while more DEGs were contributed by genome doubling in subgenome D (Figure 3A).

**Figure 3.**
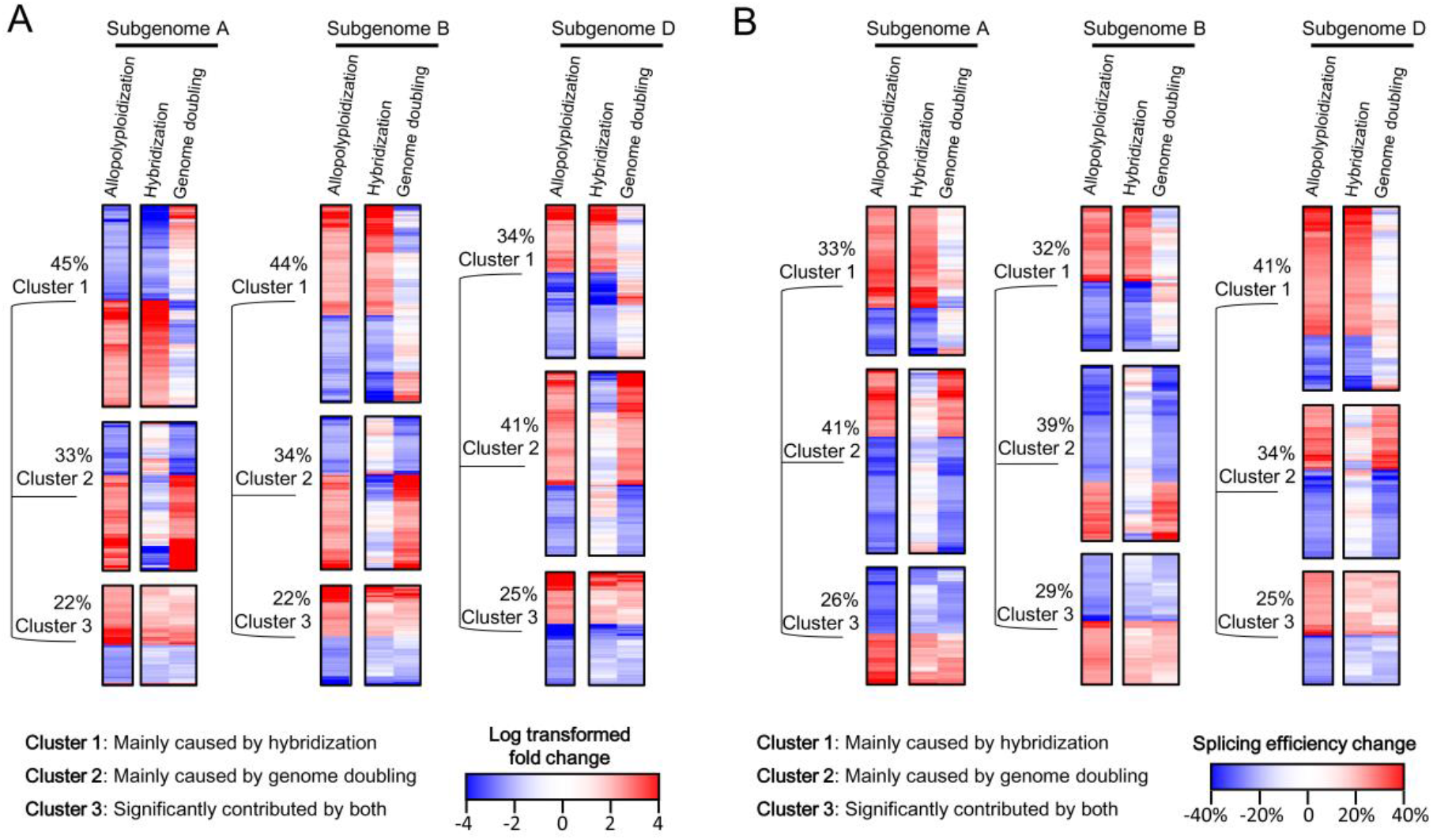
Classification of the differentially expressed genes and differentially spliced introns identified in the allopolyploids when compared with their parents. **(A)** Classification of the DEGs identified in the allopolyploid compared with the parents in wheat combination 1. All DEGs can be classified into three clusters: expression changes mainly caused by hybridization (Cluster 1), expression changes mainly caused by genome doubling (Cluster 2) and expression changes significantly contributed by both hybridization and genome doubling with the same trend (Cluster 3). The expression fold changes caused by hybridization and genome doubling, as well as their combined effect, are shown in heatmaps. Up-regulated and down-regulated genes are colored in red and blue, respectively. The color intensity reflects the magnitude of fold change. **(B)** Classification of the DSIs identified in the allopolyploid compared with the parents in wheat combination 1. All DSIs can be classified into three clusters: splicing efficiency changes mainly caused by hybridization (Cluster 1), splicing efficiency changes mainly caused by genome doubling (Cluster 2) and splicing efficiency changes significantly contributed by both hybridization and genome doubling with the same trend (Cluster 3). The splicing efficiency changes caused by hybridization and genome doubling, as well as their combined effect, are shown in heatmaps. Introns with increased and decreased splicing efficiency are colored in red and blue, respectively. The color intensity reflects the magnitude of splicing efficiency change.

It has long been considered that the heterosis in interspecific hybrids can be permanently fixed through genome doubling to form allopolyploids (Chen, 2010; Chen, 2013; Comai, 2005). The fixation of heterosis can confer advantages to allopolyploids in adaptive evolution (Chen, 2010; Comai, 2005). However, little is known about how much of the heterosis in hybrids can be fixed in allopolyploids, or exactly what role genome doubling plays in the fixation of heterosis. We found that most of the transcriptome reprogramming which occurred in hybrids cannot be fixed in allopolyploids after genome doubling (Figure 2A-2D). As transcriptome reprogramming is an important contributor to heterosis (Chen, 2010; Chen, 2013), our results suggest that most of the heterosis resulting from transcriptome reprogramming in interspecific hybrids cannot be fixed in allopolyploids due to the “recovering effect” of genome doubling.

Heterosis, or hybrid vigor, is one of the most important applications of genetics in crop breeding (Chen, 2013; Huang et al., 2016; Huang et al., 2015). Hybridization can induce dramatic transcriptome reprogramming which serves as an important source of heterosis in hybrids (Chen, 2013; Yoo et al., 2013). The transcriptome reprogramming in hybrids has typically been considered to be caused by the merging of two genomes or the “genome shock” resulting from genome merger (Chen, 2013). Our results suggested that a large proportion of the hybridization-induced transcriptome reprogramming in hybrids (Group 1 in Figure 2A and 2B) was not attributed to genome merger, as it was recovered to parental level in the allopolyploids possessing merged genomes. This proportion of transcriptome reprogramming in hybrids was probably due to other factors, such as the reduction of homologous chromosome sets in hybrids, since both parents and allopolyploids have two copies of homologous chromosome sets but hybrids only have one. In addition, several previous studies have demonstrated that hybridization-induced DNA methylation alterations can also be recovered in allopolyploids after genome doubling (Beaulieu et al., 2009; Hegarty et al., 2011; Qi et al., 2018; Xu et al., 2012). Together with our findings at gene transcriptional and splicing levels, this “recovering effect” of genome doubling implies a novel gene expression regulatory mechanism which is worth further investigation.

## Methods

### Data source, reads alignment and identification of differentially expressed genes

All the raw RNA-sequencing datasets of wheat and brassica cross combinations were published by previous studies and downloaded from NCBI SRA database (Supplemental Table 1, https://www.ncbi.nlm.nih.gov/sra/) (Hao et al., 2017; Zhang et al., 2018). In summary, two tetraploids of *T. turgidum* (AABB, 2n = 4x = 28) (ssp. *durum* cv. Langdon (LDN) and ssp. *turgidum* accession AS2255) were used as female parents, and the diploid *Ae. tauschii* accession AS60 (DD, 2n = 2x = 14) was used as male parent (Hao et al., 2017). Both LDN × AS60 (wheat combination 1) and AS2255 × AS60 (wheat combination 2) allotriploid s were produced, along with spontaneously doubled allohexaploid individuals (Hao et al., 2017). The brassica hybrid was generated by crossing *B. rapa* (A_r_A_r_, 2n = 2x = 20, as female parent) with *B. oleracea* (C_o_C_o_, 2n = 2x = 18, as male parent) (Zhang et al., 2018). The brassica tetraploid was generated by chromosome number doubling of the brassica hybrid (Zhang et al., 2018).

For wheat combinations, the reference genome sequences were download from Ensembl Plants database (https://plants.ensembl.org/, Release 44) and indexed by using *Hisat2-build* (Ver. 2.0.5) (Kim et al., 2015). RNA-Seq reads were mapped to the reference genome by using *Hisat2* (Ver. 2.0.5) with options “*-5 5 -3 5 -p 6 --no-discordant --known-splicesite-infile --reorder --no-mixed --no-unal*” (Kim et al., 2015). Only the uniquely mapped reads were retained for further analyses. Reads counts of each gene were summarized by using *HTseq-count* (Ver. 0.9.1) with options “*-f bam -r name -s no -a 0*” (Anders et al., 2015).

The reads counts were normalized by using R package *edgeR* (Ver. 3.20.9) (Robinson et al., 2010). The expression abundance was calculated as RPM (Reads assigned Per Million mapped reads). For wheat combination 2, the *p* values of each comparisons were calculated using R package *edgeR* (Ver. 3.20.9) (Robinson et al., 2010). As the *T. turgidum* cv. LDN in wheat combination 1 only has one replicate, the *p* values were calculated with Fisher’s exact test in R to eliminate the potential bias caused by different replication numbers. Raw *p* values were further adjusted by using the Benjamini & Hochbergcorrected false discovery rate (FDR). The differentially expressed genes were identified with criteria “FDR < 0.05 and fold change >=2”.

For brassica combination, the genome index was built using *Hisat2-build* (Ver. 2.0.5) with options “*--ss combined*.*ss --exon combined*.*exon*” (Kim et al., 2015). RNA-Seq reads were mapped to the reference genome by using *Hisat2* (Ver. 2.0.5) with options “*-5 5 -3 5 -p 6 --no-discordant --reorder --no-mixed --no-unal*” (Kim et al., 2015). Only the uniquely mapped reads were retained for further analyses. Reads counts of each gene were summarized by using *HTseq-count* (Ver. 0.9.1) with options “*-f bam -r name -s no -a 0*” (Anders et al., 2015). The read counts were normalized using R package *edgeR* (Ver. 3.20.9) (Robinson et al., 2010). The expression abundance was calculated as RPM. The *p* values of each comparisons were calculated with R package *edgeR* (Ver. 3.20.9) and further adjusted with Benjamini & Hochbergcorrected false discovery rate (FDR) (Robinson et al., 2010). The differentially expressed genes were identified with criteria “FDR < 0.05 and fold change >=2”.

### Splicing efficiency calculation and identification of differentially spliced introns

The splicing efficiency of an intron was defined as the expression percentage of the spliced isoforms among all the expressed isoforms (Brooks et al., 2011). To improve the accuracy of splicing efficiency quantification, introns with total reads number (spliced mapped reads and reads mapped across splice sites) less than 10 were filter out. *P* values for splicing efficiency comparisons were calculated with Cochran–Mantel–Haenszel test and further adjusted with Benjamini & Hochbergcorrected false discovery rate (FDR). Differentially spliced introns were identified with criteria “FDR < 0.05 and splicing efficiency difference >=20%”.

### Classification of differentially expressed genes

To simplify the description, gene expression fold changes (FC) calculated from Hybrid-vs-Parents, Allopolyploid-vs-Hybrid and Allopolyploid-vs-parents were referred to as FC_hybrid_, FC_genome-doubling_, and FC_allopolyploidization_, respectively. The criteria used in the classification of different groups of DEGs identified in the hybrids and allopolyploids (Figure 2A; Figure 3A; Supplemental Figure 9) were listed as below.

- Group 1 in Figure 2A 2 > FC_hybrid_ * FC_genome-doubling_ > 0.5 AND (FC_genome-doubling_ <0.67 OR FC_genome-doubling_ >1.5)
- Group 2 in Figure 2A (FC_hybrid_ >=2 AND FC_allopolyploidization_ >=2) OR (FC_hybrid_ <=0.5 AND FC_allopolyploidization_ <=0.5)
- Group 3 in Figure 2A (FC_hybrid_ >=2 AND FC_genome-doubling_ >=2) OR (FC_hybrid_ <=0.5 AND FC_genome-doubling_ <=0.5)
- Cluster 1 in Figure 3A and Supplemental Figure 9 ((FC_hybrid_ >= 2 OR FC_hybrid_ <= 0.5) AND 2 > FC_genome-doubling_ >0.5) OR ((FC_hybrid_ >= 2 AND FC_genome-doubling_ <= 0.5 AND FC_allopolyploidization_ >= 2) OR (FC_genome-doubling_ >= 2 AND FC_hybrid_ <=0.5 AND FC_allopolyploidization_ <= 0.5))
- Cluster 2 in Figure 3A and Supplemental Figure 9 ((FC_genome-doubling_ >= 2 OR FC_genome-doubling_ <= 0.5) AND 2 > FC_hybrid_ >0.5) OR ((FC_genome-doubling_ >= 2 and FC_hybrid_ <= 0.5 AND FC_allopolyploidization_ >= 2) OR (FC_hybrid_ >= 2 AND FC_genome-doubling_ <=0.5 AND FC_allopolyploidization_ <= 0.5))
- Cluster 3 in Figure 3A and Supplemental Figure 9 (FC_hybrid_ >=2 AND FC_genome-doubling_ >=2) OR (FC_hybrid_ <=0.5 AND FC_genome-doubling_ <=0.5)

### Classification of differentially spliced introns

To simplify the description, splicing efficiency changes (SEC) calculated from Hybrid-vs-Parents, Allopolyploid-vs-Hybrid and Allopolyploid-vs-parents were referred to as SEC_hybrid_, SEC_genome-doubling_, and SEC_allopolyploidization_, respectively. The criteria used in the classification of different groups of DSIs identified in the hybrids and allopolyploids (Figure 2B; Figure 3B; Supplemental Figure 10) were listed as below.

- Group 1 in Figure 2B (|SEC_hybrid_ + SEC_genome-doubling_ | < 0.2) AND (|SEC_genome-doubling_| > 0.1)
- Group 2 in Figure 2B (SEC_hybrid_ >=0.2 AND SEC_allopolyploidization_ >=0.2) OR (SEC_hybrid_ <=−0.2 AND SEC_allopolyploidization_ <=−0.2)
- Group 3 in Figure 2B (SEC_hybrid_ >=0.2 AND SEC_genome-doubling_ >=0.2) OR (SEC_hybrid_ <=−0.2 AND SEC_genome-doubling_ <=−0.2)
- Cluster 1 in Figure 3B and Supplemental Figure 10 (|SEC_hybrid_| >= 0.2 AND |SEC_genome-doubling_| <0.2) OR ((SEC_hybrid_ >=0.2 AND SEC_genome-doubling_ <= −0.2 AND SEC_allopolyploidization_ >= 0.2) OR (SEC_genome-doubling_ >= 0.2 AND SEC_hybrid_ <=−0.2 AND SEC_allopolyploidization_ <= −0.2))
- Cluster 2 in Figure 3B and Supplemental Figure 10 (|SEC_genome-doubling_| >= 0.2 AND |SEC_hybrid_| <0.2) OR ((SEC_genome-doubling_ >= 0.2 AND SEC_hybrid_ <= −0.2 AND SEC_allopolyploidization_ >= 0.2) OR (SEC_hybrid_ >= 0.2 AND SEC_genome-doubling_ <=−0.2 AND SEC_allopolyploidization_ <= −0.2))
- Cluster 3 in Figure 3B and Supplemental Figure 10 (SEC_hybrid_ >=0.2 AND SEC_genome-doubling_ >=0.2) OR (SEC_hybrid_ <=−0.2 AND SEC_genome-doubling_ <=−0.2)

## Author Contributions

Z.L., Q.S., and Z.N. conceived the research; Z.L. and J.Q. designed the analyses; Z.L., J.Q., R.M. and H.L. performed the data analysis; J.Q. and Z.L. wrote the manuscript with input from all authors.

## Acknowledgments

This work was financially supported by the National Natural Science Foundation of China (Grant No. 31601304, 31601305), Shaanxi Natural Science Foundation (Grant No. 2017JQ3023) and the Basic Scientific Research Foundation of Northwest A&F University (Grant No. Z109021806, Z1090220096). No conflict of interest declared.

## Notes

### Competing Interest Statement

The authors have declared no competing interest.

